# The C-terminal actin binding domain of talin forms an asymmetric catch bond with F-actin

**DOI:** 10.1101/2020.09.01.276568

**Authors:** Leanna M. Owen, Nick A. Bax, William I. Weis, Alexander R. Dunn

**Author notes:** Corresponding authors: (W.I.W.); (A.R.D.).

## Abstract

Focal adhesions (FAs) are large, integrin-based adhesion complexes that link cells to the extracellular matrix (ECM). Previous work demonstrates that FAs form only when and where they are necessary to transmit force between the cellular cytoskeleton and the ECM, but how this occurs remains poorly understood. Talin is a 270 kDa adapter protein that links integrins to filamentous (F)-actin and recruits additional components during FA assembly in a force-dependent manner. Cell biological and developmental data demonstrate that the third, and C-terminal, F-actin binding site (ABS3) of talin is required for normal FA formation. However, ABS3 binds F-actin only weakly in *in vitro*, biochemical assays. We used a single-molecule optical trap assay to examine how and whether ABS3 binds F-actin under physiologically relevant, pN mechanical loads. We find that ABS3 forms a directional catch bond with F-actin when force is applied towards the pointed end of the actin filament, with binding lifetimes more than 100-fold longer than when force is applied towards the barbed end. Long-lived bonds to F-actin under load require the ABS3 C-terminal dimerization domain, whose cleavage is known to regulate focal adhesion turnover. Our results support a mechanism in which talin ABS3 preferentially binds and orients actin filaments with barbed ends facing the cell periphery, thus nucleating long-range order in the actin cytoskeleton. We suggest that talin ABS3 may function as a molecular AND gate that allows FA growth only when sufficient integrin density, F-actin polarization, and mechanical tension are simultaneously present.

## Main Text

Precise and dynamically regulated force transmission between cells and the extracellular matrix (ECM) is a requirement for cell migration, tissue repair, and more broadly the construction of multicellular animal life. Adhesion to the ECM is mediated in large part by integrins, a family of heterodimeric transmembrane proteins that assemble into large adhesion complexes, here referred to generically as focal adhesions (FAs). FA assembly is precisely controlled, such that FAs form only when and where they are necessary to transmit force between the cytoskeleton and the ECM. However, the mechanism by which mechanical force regulates FA nucleation and growth remains poorly understood.

The connection between integrins and the actin cytoskeleton is mediated in large part by talin, a 270 kDa scaffolding protein that directly binds integrins, F-actin, and other FA components (Klapholz and Brown, 2017). Talin possesses three F-actin binding sites, termed ABS1-3. In addition, the talin rod domain is constructed of a series helical bundle domains that unfold under loads of 5-25 pN (del Rio *et al*., 2009; Yao *et al*., 2014, 2016) to expose at least 9 binding sites for the F-actin binding protein vinculin (Gingras *et al*., 2005), which is thought to reinforce the adhesion.

Despite the multiple direct and indirect (through vinculin) interactions of talin with F-actin, binding by the C-terminal ABS3 to F-actin is essential for its biological functions (Goult, Yan and Schwartz, 2018). ABS3 consists of a five-helix bundle along with an isolated C-terminal alpha helix that mediates homodimerization (Smith and McCann, 2007; Goult *et al*., 2008; Bate *et al*., 2012). ABS3 mutations that disrupt F-actin binding hinder the initial stages of adhesion formation (Giannone *et al*., 2003; Jiang *et al*., 2003), as well as subsequent FA assembly (Giannone *et al*., 2003; Kopp *et al*., 2010; Atherton *et al*., 2015) and cell spreading (Kopp *et al*., 2010). *Drosophila* with ABS3 mutations that disrupt F-actin binding show defects in germband retraction resembling talin-null embryos, as well as defects in embryonic muscle-tendon attachment and epithelial sheet adhesion in the adult wing (Franco-Cea *et al*., 2010; Klapholz *et al*., 2015). Despite its functional importance, ABS3 binds F-actin only weakly in co-sedimentation assays (Senetar, Foster and McCann, 2004; Gingras *et al*., 2008; Srivastava *et al*., 2008). A pioneering study in live cells was interpreted to indicate that talin forms a weak, 2 pN slip bond to F-actin that required ABS3 (Jiang *et al*., 2003). However, to our knowledge it is not known how, or even whether, mechanical load may affect the stability of the ABS3–F-actin bond.

We used a previously described single-molecule assay (Buckley *et al*., 2014; Huang *et al*., 2017) to characterize the binding of human ABS3 (residues 2293-2541) to F-actin (**Fig. 1Ai, Bi**). We first characterized this interaction in the absence of externally applied load and observed only short ABS3-F-actin bond lifetimes with a mean of 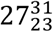 ms (95% confidence interval; CI) **(Fig. Aiii**). In contrast, in the presence of biologically relevant loads (Ringer *et al*., 2017) (**Fig. 1Bi**) we observed stable binding of ABS3 to actin, with bond lifetimes in the tens and sometimes hundreds of seconds (**Fig. 1Bii**).

**Figure 1:**
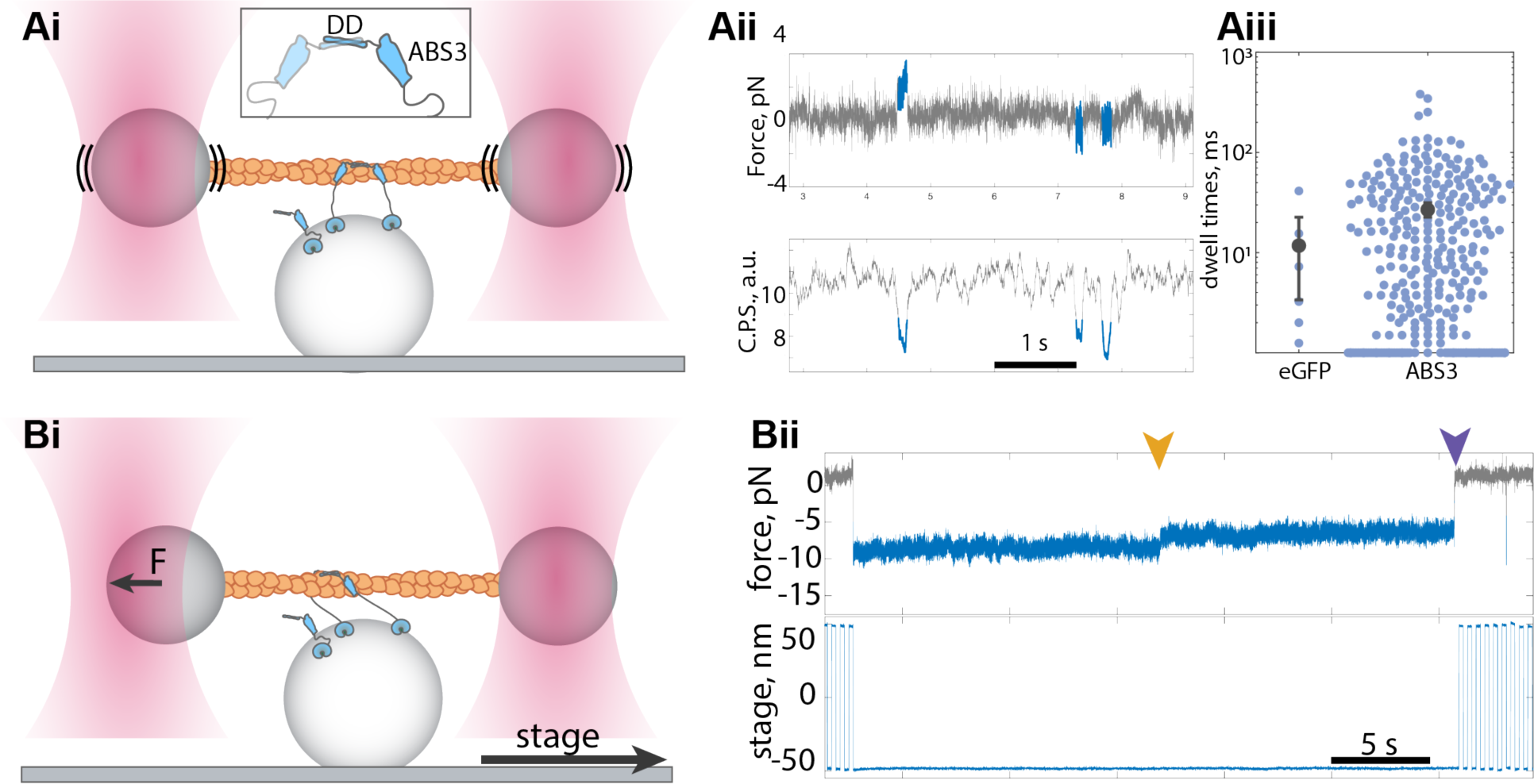
Talin-1 ABS3 binding to actin is stabilized by force. (**Ai**) No-load three bead optical trap experimental setup. ABS3 (light blue) is immobilized via an N-terminal HaloTag domain on a microscope coverslip sparsely decorated with 1.5 μm diameter platform beads. A single actin filament (orange) is stretched between two optically trapped 1 μm diameter beads. The microscope stage is positioned such that a platform bead is positioned directly underneath the actin filament. (**Aii**) Sample trace of the position of the trapped beads to detect binding of ABS3 to F-actin. In the absence of applied load, ABS3 binding to F-actin can be detected by a damping of high-frequency Brownian fluctuations (cumulative power spectrum, C.P.S. > 0.3-10 kHz) in the trapped beads. (**Aiii**) Binding lifetimes detected with this method. N=291 binding events detected from sampling the ABS3 construct from 1,419 seconds of recorded data. N=8 binding events detected from sampling an eGFP-coated surface from 2,188 seconds. (**Bi**) Loaded three bead optical trap experimental setup. Same as **Ai**, except the stage is translated parallel to the actin filament in a trapezoidal wave form. (**Bii**) If one or more ABS3 molecules on the platform bead bind to the actin filament while the stage is moving, one of the trapped beads is pulled out of its equilibrium position (demarcated by the blue force trace), thus applying load to the ABS3-F-actin bond. When this occurs the stage motion is halted until all of the protein complexes unbind and the trapped beads return to their equilibrium positions (marked with purple notch, gray trace). Reported lifetimes are last steps of binding events, which for this sample trace is the time between yellow and purple arrows, or the full lifetime of single-step events.

Strikingly, we also observed that nearly all long (>0.5 s) binding events associated with a single actin filament were associated with load on only one of the two traps (**Fig. 2A**). This consistency in the preferred direction of long-lived binding events was observed for a given actin filament across multiple platforms, indicating that this behavior did not reflect ABS3 orientation on the surface. When we reversed the orientation of the actin filament by switching the position of the two beads in the optical traps the preferred loading direction also switched (data not shown). This result strongly suggested that the relationship between the polarity of the actin filament and the vector of applied force regulated the stability of the ABS3 to F-actin bond.

**Figure 2:**
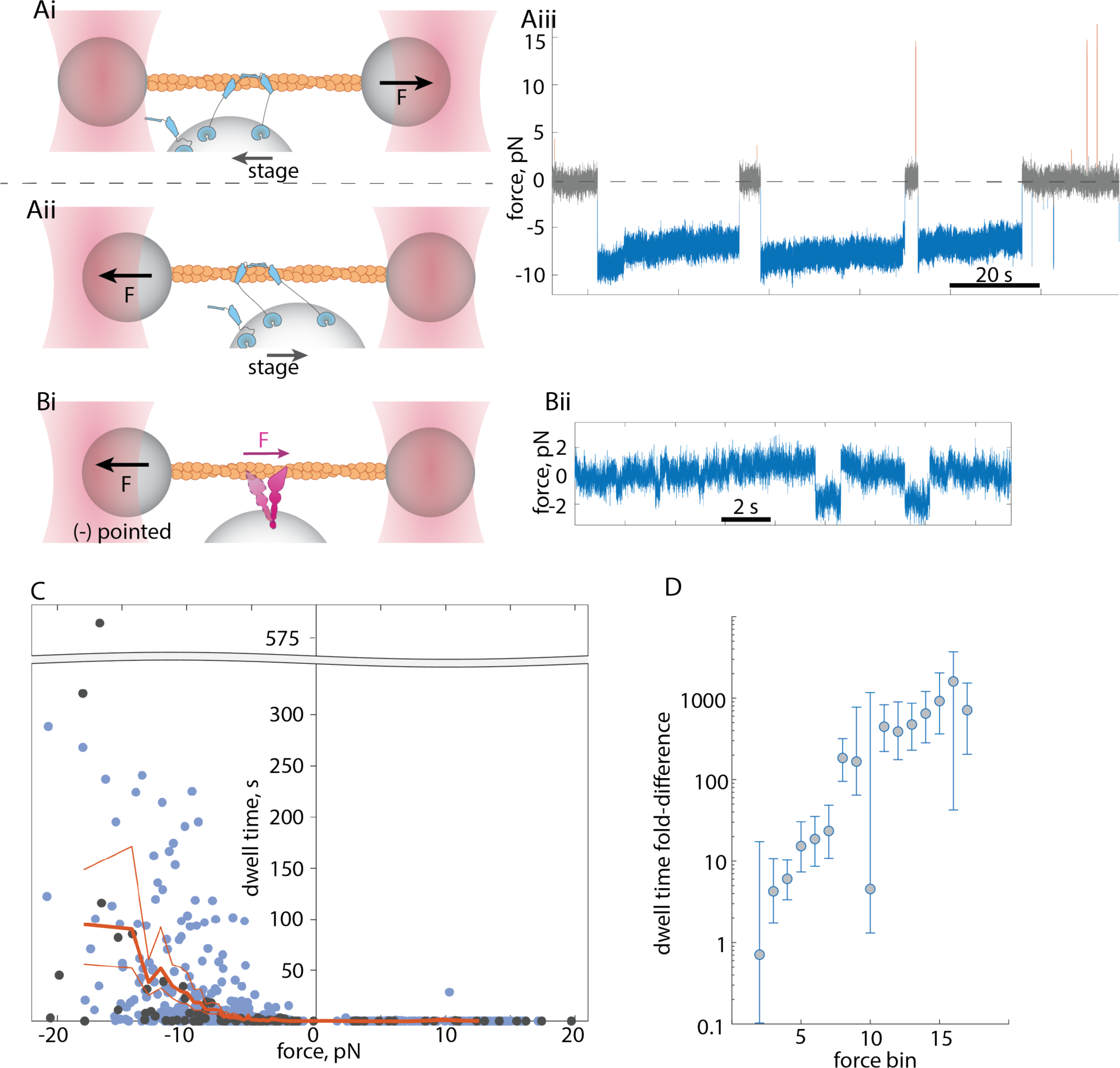
Talin-1 forms a catch bond with F-actin when force is applied towards the pointed end of the actin filament. (**A**) An asymmetry in bond lifetimes is observed for a single actin filament. (**Ai, Aii**) Load on the *right* (**Ai**) or *left* (**Aii**) trapped bead. (**Aiii**) Sample force trace. Load on the right bead is indicated in orange, while load on the left bead is shown in blue. (**B**) (**Bi**) A myosin VI construct steps along actin filaments, pulling the bead attached to the (-) end of the filament out of the trap. (**Bii**) Sample myosin trace. In this example, the left bead is attached to the F-actin (-) end. (**C**) ABS3 binding dwell times measured as a function of force. Forces directed towards the pointed (-) end of the filament correspond to negative force values and forces directed towards the barbed (+) end correspond to positive force values. *Dark gray*: binding lifetimes of events measured in the directionality assay in which the actin filament polarity was explicitly measured for each dumbbell (N=2 flow cells, N=3 dumbbells, N=47 events). *Lavender*: binding dwell times for which the actin filament polarity was statistically inferred. *Orange*: mean dwell times (20 data points per bin), with 95% confidence intervals shown in thinner lines (N = 777 binding events, N=42 dumbbells, N=18 flow cells). (**D**) Fold-difference in bond lifetime for forces applied towards the pointed vs. barbed end of the actin filament, with 95% confidence intervals per 1 pN bin.

We next determined whether long-lived binding events corresponded to load oriented towards the F-actin barbed (+) or pointed (-) end by using a previously reported assay in which the polarity of a given actin filament is determined using the activity of a myosin VI construct (**Fig. 2B**) (Huang *et al*., 2017). (Myosin VI is a nonmuscle myosin that moves toward the (-) end of F-actin.) ABS3 binding events longer than 0.48 s only occurred when force was applied towards the F-actin pointed (-) end (**Fig. 2C**). This asymmetry was so striking that we could infer the polarity of actin filaments in previous datasets with high confidence, and in so doing quantify force-binding lifetime relationships relative to actin filament polarity (**Fig. 2C**). The difference in binding lifetimes was most striking for forces above 10 pN, where the difference in lifetimes was >100-fold (**Fig. 2D**). Only 125/777 (17%) of the detectable binding events occurred under (+)-end directed loads, suggesting that the measured asymmetry in lifetimes constitutes a lower limit.

The C-terminal dimerization domain of ABS3 is required for actin cosedimentation and for the function of ABS3 *in vivo* (Gingras *et al*., 2008; Franco-Cea *et al*., 2010; Bate *et al*., 2012; Klapholz *et al*., 2015). To determine how this domain contributes to the F-actin binding, we created a construct containing talin residues 2293-2493 that lacked the C-terminal dimerization domain, termed ABS3ΔDD (**Fig. 3A**). Consistent with earlier reports, this construct cosedimented with F-actin weakly (data not shown) (Gingras *et al*., 2008). Moreover, we detected no binding events in the optical trap assay longer than 25 ms using the low-force assay (N=11 events detected in 956 s, mean 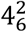 ms), and no binding under load that could be differentiated from negative controls.

**Figure 3:**
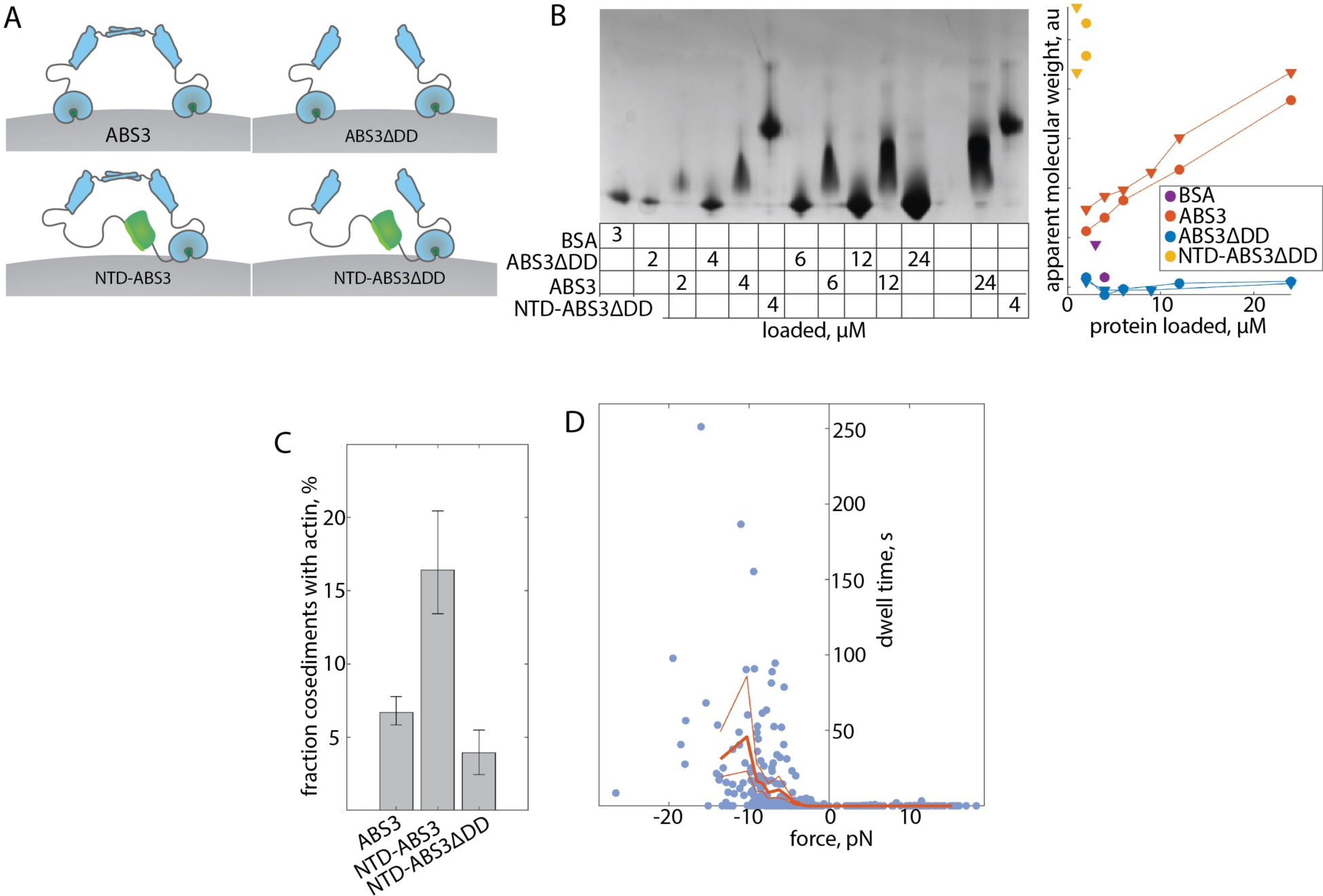
The C-terminal dimerization domain (DD) is required for F-actin binding under load. (**A**) The constructs used in this study (see text). (**B**) Native PAGE of BSA (MW=66.5 as monomer), ABS3 (64.2 kDa if monomeric), ABS3ΔDD (58.5 kDa, monomer) and NTD-ABS3ΔDD (113 kDa). Note the gradual shift in apparent molecular weight as a function of ABS3 concentration, suggestive of a dynamic equilibrium of dimerization. • and ▾denote distances traveled of peaks in different replicates. (C) Actin cosedimentation of 4 uM ABS3, 2 uM NTD-ABS3, or 2 uM NTD-ABS3ΔDD with 20 µM actin. N=3 replicates. (D) Force-lifetime plots of binding events detected for the NTD-ABS3 construct (lavender) with mean lifetime and confidence intervals on the mean plotted in orange (20 data points per bin). N=386 binding events, N=18 dumbbells, N=8 flow cells.

The lack of detectable binding by ABS3ΔDD in the optical trap assay suggested two non-exclusive possibilities for the role of the dimerization domain: the dimerization domain is required for strong binding of a single ABS3 to F-actin, and/or the recruitment of two ABS3 domains is required for strong binding. To distinguish among these possibilities, we first more closely examined the dimerization of ABS3, which had been reported to form a constitutive dimer (Azizi *et al*., 2020). In our hands, ABS3 dimerization is a dynamic equilibrium at micromolar concentrations, as determined by native PAGE (**Fig. 3B**).

Next, we created two constructs, termed <u>N</u>-terminal <u>d</u>imer (NTD)-ABS3 and NTD-ABS3ΔDD, in which split GFP (Cabantous, Terwilliger and Waldo, 2005) was used to make stable dimers of either ABS3 or ABS3ΔDD (**Fig. 3A**). Although NTD-ABS3ΔDD retained F-actin affinity in the co-sedimentation assay (**Fig. 3C**), it showed low binding activity in the optical trap assay, with no events longer than 0.8 s in the presence of applied force. Thus, dimerization alone was not sufficient to rescue long-lived binding in the presence of load. For NTD-ABS3, forced N-terminal dimerization modestly enhanced F-actin co-sedimentation (**Fig. 3C**). However, in the optical trap assay, F-actin binding lifetimes for NTD-ABS3 were similar to those of ABS3 both in the absence (N=510 events detected in 695 s, mean binding lifetime 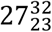 ms; 95% CI) or presence of load (**Fig. 3D**). This degree of similarity strongly suggests that ABS3 also forms stable, load-bearing bonds to F-actin only when its C-terminal actin binding domain is present. Although it must be interpreted with care, we also did not observe substeps for the dissociation of ABS3 or NTD-ABS3 from F-actin, which if present would provide evidence for sequential unbinding of individual ABS3 domains. In total, a reasonable interpretation is that *i*) definitively monomeric ABS3 (ABS3-ΔDD) binds F-actin only transiently under load, *ii*) the DD is required for stable F-actin binding under load, and *iii*) each dissociation step we observe reflects the coordinated detachment, within our temporal resolution, of an ABS3 dimer from F-actin.

The force sensitivity and directionality of the interaction of ABS3 with F-actin is remarkable. We had previously reported that the talin binding partner vinculin also forms a directional catch bond with F-actin (Huang *et al*., 2017): like talin, the bond between vinculin and F-actin is stabilized when load is oriented toward the pointed end of the actin filament. However, both the force dependence and directional preference of ABS3 are extreme – the mean measured lifetimes and the actin polarity preference each increase more than 100-fold under load, enhancements that are ∼10-fold larger than those observed for vinculin (Huang *et al*., 2017).

Indirect evidence suggests that the force and directional sensitivities of vinculin and talin ABS3 may act in concert: Knockout of vinculin worsens ABS3 mutant phenotypes in mammalian cells in culture and in *Drosophila* (Klapholz *et al*., 2015), and expression of constitutively active vinculin can compensate for ABS3 disruption in cell culture (Atherton *et al*., 2015). Whether the talin ABS1 and ABS2 domains, and more broadly the many other F-actin binding proteins present at FAs, show similar force- and/or directional sensitivity is not known. It is also possible that force- and direction-sensitive actin-binding proteins, including vinculin, help to polarize F-actin at cell-ECM adhesions. These and related questions present interesting opportunities for future investigations.

How exactly cells regulate the number, size and placement of FAs has remained unclear. As a partial answer, we speculate that ABS3 may function as a molecular AND gate that limits the formation of FAs to highly specific circumstances (**Fig. 4**). As suggested previously, it is plausible that integrins must be of sufficient local areal density to anchor talin dimers at both N-termini (Roca-Cusachs *et al*., 2009; Schvartzman *et al*., 2011). Further, our results show that substantial, >5 pN loads are required for ABS3 to form long-lived bonds to F-actin. This condition requires that load-bearing connections between the ECM, integrin, and talin are intact, and that the proximal ECM is sufficiently stiff to resist cell-generated forces. Finally, load must be oriented toward the F-actin pointed end. This stabilizes oriented F-actin that is under load due to either barbed-end polymerization forces or tension generated by nonmuscle myosin II, and allows mis-oriented F-actin to escape. Simulation suggests that the resulting polarization of F-actin can potentially drive cellular-scale organization of the actin cytoskeleton to generate efficient traction and produce directional cell migration (Thievessen *et al*., 2015; Rahman *et al*., 2016; Huang *et al*., 2017).

**Figure 4.**
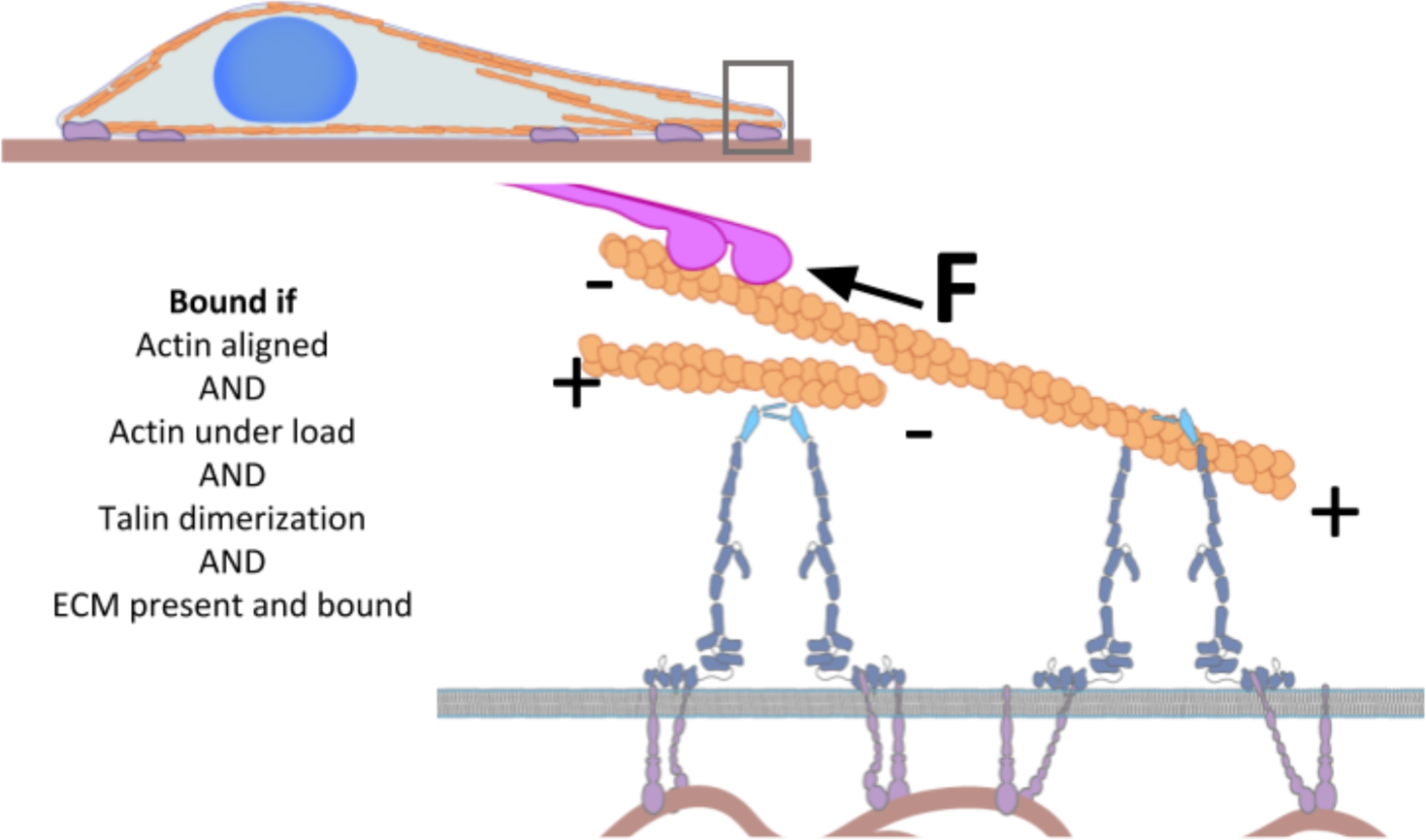
Talin ABS3 functions as a molecular AND gate (see text). Integrin cartoon modified from Zhu *et al*., 2017.

In summary, we propose that the F-actin binding properties of ABS3 have evolved to limit FA formation precisely to when and where a force-transmitting anchorage between the cytoskeleton and the ECM is required. Further work is required to test this hypothesis with *in vivo* experiments, and to test whether a similar form of autoregulation may apply to other cellular adhesion complexes.

## Materials and Methods

### Constructs

The HaloTag-ABS3 expression vector was constructed by inserting the DNA encoding the Halo Tag, a linker sequence, and the human talin-1 C-terminal actin binding domain (amino acids 2293-2541) into the pPROEX HTa expression vector to generate an in-frame fusion consisting of a 6xHis-tag, HaloTag, 12 residue linker (SGLGGSGGGGSG), and ABS3. The GFP11-ABS3 vector was generated by insertion of the sequence for GFP11 and ABS3 into a pD444-NH: T5-His-ORF, Ecoli-Elec D expression vector to create an in-frame fusion of His6-tag, a (GGSGG)x2 linker, GFP11, (GGSGG)x6, and ABS3. The GFP 1-10 OPT-HaloTag-ABS3 expression vector was generated by insertion of the sequence for GFP 1-10 OPT, HaloTag, and ABS3 into pD444-NH: T5-His-ORF, Ecoli-Elec D to create an in-frame fusion of a 6xHis-tag, GFP 1-10 OPT, (GGSGG)x2, HaloTag, (GGSGG)x4, and ABS3. The HaloTag-ABS3ΔDD expression vector was generated from the HaloTag-ABS3 expression vector by the insertion of two stop codons before amino acid M2494 of the original talin-1 gene. The GFP11-ABS3ΔDD and GFP 1-10 OPT-HaloTag-ABS3ΔDD expression vectors were generated by site-directed mutagenesis of amino acid M2494 of the original talin-1 gene to generate a stop codon in its place.

### Protein preparation

The GFP11 and GFP 1-10 OPT split GFP constructs were co-expressed in BL21 cultures to facilitate split GFP formation and stabilization (Cabantous, Terwilliger and Waldo, 2005). All recombinant fusion proteins were expressed in BL21 (DE3) E. coli cells grown in 1-2 L LB media at 37 °C with 50 μg/mL kanamycin and 50 μg/mL carbenicillin to an optical density of 0.5-0.8, then induced with 0.5-1 mM IPTG. After induction, cells were grown for 4 hours at 37 °C. Cells were centrifuged for 20 minutes at 5000 x g. Cell pellets were resuspended in approximately 50 mL of 20 mM Tris pH 8.0, 50 mM KCl and stored at - 80 °C until protein purification.

Following treatment with DNAse, lysozyme, and 3 cOmplete, EDTA-free Protease Inhibitor Cocktail tablets (Roche), cells were lysed with an Emulsiflex (Avastin) cell disrupter and the lysate was centrifuged at 4 °C 12,000 x g for 30 minutes. β-mercaptoethanol was added to 2 mM. Clarified lysate was incubated with 10 mL of Ni-NTA resin for 2 hours on a rocker at 4 °C. Protein was washed with 3 bed volumes of wash buffer (300 mM NaCl, 50 mM NaH_2_PO_4_, 20 mM imidazole, pH 7.4) and eluted in one bed volume of elution buffer (300 mM NaCl, 50 mM NaH_2_PO_4_, 250 mM imidazole, pH 7.4). The eluate was passed through a 0.45 μm PES syringe filter (Millipore) and diluted to a final volume of 50 mL in 20 mM Tris pH 8.0. The filtered eluate was applied to a MonoQ anion exchange column (GE) in 20 mM Tris pH 8.0 buffer and run with a 0-1 M KCl gradient. The proteins were further purified by Superdex 200 (GE) gel filtration chromatography in 20 mM Tris, pH 8.0, 50 mM KCl, 1 mM DTT. Protein was concentrated on a 30kDa MWCO Amicon Ultra centrifugal filter to between 7.7 and 56 μM, as measured by absorbance at 280 nm, and flash-frozen for storage at -80 °C.

eGFP was purified and labeled with HaloTag Succinimidyl Ester (O4) Ligand (Promega, P6751) as previously described (Huang, D.L., *et al*. 2017). Ligand labeling densities > 1 Halo Tag ligand per eGFP were required for activity of the ABS3 construct.

### Actin biotinylation protocol

F-actin used in the optical trap assay was purified from rabbit skeletal muscle, as previously described (Spudich and Watt, 1971) then flash frozen and at -80 °C in G-Actin buffer (5 mM Tris-HCl pH 8.0, 0.2 mM CaCl_2_, and 0.2 mM ATP). G-actin at 2 mg/ml was thawed in a gloved hand and stored on ice for 30 minutes, then centrifuged to pellet any aggregated protein. All centrifugation steps were carried out in a TLA-120 rotor (Beckman) at 60k RPM and 4 °C. for 20 minutes. 10x F-actin buffer pH 7.5 (10x FB7.5: 500 mM KCl, 200 mM Tris pH 7.5, 20 mM MgCl_2_, 2 mM CaCl_2_, 10 mM DTT) was added to the supernatant and the G-actin was allowed to polymerize for at least 1 hour at room temperature. The polymerized F-actin was pelleted by centrifugation. The supernatant was aspirated, taking care not to disturb the pellet and washed 3 times with conjugation buffer (Conjugation buffer:20 mM Phosphate Buffer pH 8, 50 mM KCl, 2 mM MgCl_2_, 0.2 mM CaCl_2_, 1 mM ATP, 1 mM DTT). The pellet was then resuspended in conjugation buffer containing 1 mM Biotin-NHS (Sigma 203118) by pipetting up and down. The ratio of biotin-NHS to actin monomers was approximately 20:1. After 30 minutes, the conjugation reaction was quenched by adding 1M glycine to a final concentration of 1 mM glycine. The conjugated actin was pelleted by centrifugation. The pellet was resuspended in G-actin buffer and dialyzed against G-actin buffer with 1 mM DTT in a 10k MWCO Slide-A-Lyzer dialysis cassette overnight with three buffer exchanges. After dialysis, the biotinylated G-actin was centrifuged to remove any aggregates. The concentration of G-Actin in the supernatant was measured using OD at 290 nm (to avoid the prominent absorption peak from ATP in the buffer) and found to be approximately 1 mg/mL. The supernatant was flash frozen in single use aliquots and stored at -80 °C.

### Flow cell preparation

Nitrocellulose-coated coverslips with attached platform beads and flow chambers were prepared as described previously (Huang, D.L, et al. 2017).

Halo tagged ABS3 fusion constructs were attached to the coverslip and platform bead surface via HaloLigand-eGFP nonspecifically adsorbed onto the coverslip and platform bead surfaces.

ABS3 constructs and HaloLigand-eGFP were diluted in F-actin buffer at pH 7.5 (FB7.5: 50 mM KCl, 20 mM HEPES pH 7.5, 2 mM MgCl_2_, 0.2 mM CaCl_2_, 1 mM DTT). Other dilutions were done in F-buffer at pH 7.1 (FB7.1: 50 mM KCl, 20 mM HEPES pH 7.1, 2 mM MgCl_2_, 0.2 mM CaCl_2_, 1 mM DTT, 1 mM ATP). ATP and DTT were added on the day of the experiment, shortly before flow cell preparation began.

Streptavadin-coated beads (Bangs Laboratories, Inc; CP01004) were freshly washed each day by centrifuging for 1 minute at 5000 x *g* and then resuspending in 20 mM HEPES pH 6.8, 50 mM KCl, 1 mM DTT, 1 mg/mL BSA at least 4 times with 5 min bath sonication between wash steps. Beads were then diluted at least 50-fold in 20 mM Tris pH 8, 50 mM KCl, 1 mg/mL BSA prior to their dilution into the trapping solution (see below).

Flow cells were prepared as followed:

All injections were 10 uL.

1. FB7.5 was injected into the channel.
2. HaloLigand-eGFP was injected into the channel and incubated for 2 minutes.
3. FB7.5 was injected into the channel to wash out the HaloLigand-eGFP.
4. 2% Pluronic F-127 (Sigma, P2443) in 5 mM Tris pH 8, 50 mM KCl, 2 mM MgCl_2_ was injected into the channel and incubated for 10 minutes.
5. FB7.5 was injected into the channel twice to wash out the pluronic.
6. HaloTag-ABS3, HaloTag-ABS3ΔDD, sGFP-HaloTag-ABS3×2, or sGFP-HaloTag-ABS3ΔDDx2 was injected into the channel at a concentration of 1μM, 1 μM, 50 nM-500 nM, or 500 nM (respectively) and incubated for 2 minutes.
7. FB7.5 was injected into the channel to wash out any ABS3 construct not bound to the platform beads.
8. BSA (1 mg/mL, MCLAB UBSA-100) diluted in FB7.1 was incubated in the chamber for 5 minutes.
9. The channel was then filled with a final trapping solution of 1mg/mL BSA, 0.8% glucose, 2.7 kU/mL catalase (Sigma, C40-100mg), 7.5 U/mL pyranose oxidase (Sigma P4234-250UN), 1 mM Trolox (Fisher Scientific, AC218940010), washed streptavidin-coated polystyrene microspheres (diluted 1:400 from stock concentration), and 0.2 nM biotinylated actin filaments, labeled with rhodamine-phalloidin (Cytoskeleton PHDR1), and 20 μM phalloidin in FB 7.1.

Flow cells for determining actin filament polarity and preferred direction of ABS3 binding were prepared as described previously (Huang *et al*., 2017).

### Optical Trap instrument

The optical trap used was essentially the same as described previously (Sung *et al*., 2010; Buckley *et al*., 2014; Huang *et al*., 2017). The trap was operated at a stiffness of 0.15 to 0.20 pN/nm.

### Loaded optical trap assay

The flow cell was prepared as described above and mounted on the stage of the optical trap. Streptavadin-coated beads were captured in each of the two optical trap beams, and the stage was steered to tether a single actin filament to the beads at either end to form a dumbbell. The streptavadin beads were moved apart until the actin filament was taut, with approximately 1-2 pN of tension across the filament. The actin filament was brought into close proximity of a platform bead. The stage was then oscillated parallel to the actin filament with a 20-50 nm amplitude, 10 nm/ms rise/fall rate, and a pause of 150 ms to check for displacement of either trapped bead. If displacement was detected, then the stage paused until both beads relaxed to their starting positions before restarting the oscillation.

### Analysis of loaded optical trap assay

Trap stiffness calibration was performed as has previously described (Huang *et al*., 2017).

Bead position data were down sampled to 1 kHz, and binding events were picked with custom software (MATLAB) and verified manually. The force associated with each binding event was calculated as described previously (Huang *et al*., 2017). Both single step events, which dominated this dataset, and the last steps of multi-step binding events were plotted and included in analysis. Binding events less than 20 ms were excluded from all analyses as these were also observed in a minority of negative controls.

### Assignment of actin filament polarity

Actin filament polarity was explicitly determined as described in Huang *et al*., 2017 for a subset of the data in which flow cells were prepared with a myosin VI construct consisting of porcine myosin VI (residues 1– 817) and *Archaeoglobus fulgidus* L7Ae (residues 9–118) with a C-terminal eYFP (Omabegho *et al*., 2018) on one half of the flow cell (Huang *et al*., 2017). In this case, actin filaments were held above multiple myosin VI platforms and the direction of myosin force generation was observed. The actin dumbbell was then moved to a location in the flow cell coated with ABS3, and the loaded optical trap assay was performed. Myosin stepping was observed to displace actin filaments in the same direction as strong ABS3 binding, indicating that ABS3 binds stably when force is applied towards the pointed end of the actin filament for dumbbells measured with this assay.

Actin filament polarity was inferred for the remainder of the data in which no myosin VI was present. Filament polarity could be reliably determined for dumbbells for which we measured at least two binding events longer than 0.5 seconds. In these cases, actin filament polarity was assigned such that the longer mean lifetimes were associated with pointed-end directed forces. Traces corresponding to actin dumbbells that yielded fewer than 2 long (>0.5 s) events were excluded from analysis for both ABS3 and NTD-ABS3 in the loaded assay, as the orientation of these filaments could not be clearly determined.

### No-load optical trap assay

For the ABS3 and NTD-ABS3 constructs, the flow cell was prepared as for a loaded optical trap assay. A dumbbell was formed as described above, and platform bead positions were assayed for specific binding activity with load. If binding was detected, the last binding event was allowed to terminate and the stage oscillation was turned off. Without moving the dumbbell or stage, bead displacement data was then collected at a 40 kHz sampling rate.

Because ABS3ΔDD construct and eGFP negative control produced no obvious specific binding in the presence of force, several bead positions in each of several flow cells were assayed at random.

Binding events were annotated from the above dataset as follows: The 40 kHz signal for each bead position was downsampled to 4 kHz, and a Fourier transform of a 256 data point window was taken at every point. The power of the signal above 300 Hz was calculated for each point and used to determine binding. Deviations of the summed (across the two beads) high-frequency power below 82% of mean were annotated as a binding event. Additionally, if the high frequency power of either bead dipped below 70% of that bead’s mean, that was also annotated as a binding event. ‘Holes’ between initially detected binding events of less than 4 ms were filled in such that adjacent binding events were combined.

### Actin co-sedimentation assay

Rabbit skeletal actin was polymerized at 36 μM by adding 10x F-actin buffer (500 mM KCl, 20 mM MgCl_2_, and 10 mM ATP) at room temperature for 1 hour. F-actin was stored at 4 °C and used within 3 days. 20 mM Tris pH 8, 50 mM KCl, and 1 mM DTT was added such that the final concentration of Tris, KCl, and DTT would be constant in all samples following the addition of protein. HEPES pH 6.5 was added to each sample to adjust the pH to 7.0. Purified ABS3 protein constructs were thawed and mixed with F-actin to a final concentration of 4 μM ABS3, 2 μM NTD-ABS3 or 2 μM NTD-ABS3ΔDD, such that the total concentration of actin binding domains was constant between conditions. The final concentration of buffers and salts was 20 mM HEPES, 6.8 mM Tris, 50 mM KCl, 1.1 mM MgCl_2_, 0.11 mM CaCl_2_. The pH was checked for each sample with a micro pH probe and verified to be 6.9-7.0. Samples were incubated at room temperature for 40 min to 1 hour, then centrifuged at 22 °C at 100,000 x g for 15 minutes. Supernatant and actin pellet were separated and analyzed via reducing SDS-PAGE.

### Native PAGE assay

Native PAGE was performed using NuPAGE™ 12%, Bis-Tris gels (Invitrogen). Talin ABS proteins were desalted in a 40kDa MWCO Zeba desalting column (Thermo Scientific). Proteins were prepared at the specified concentration (Fig. 3b) by mixing with 2.5 μL NativePAGE Sample Buffer (Thermo Scientific, BN2003), 10 mM DTT, and water added to a total volume of 10 μL. Samples were incubated for 15 minutes prior to loading the gel. The gel was run following manufacturer instructions with “light blue” cathode buffer (0.5% NativePAGE™ Cathode Buffer Additive 20x, 5% NativePAGE™ Running Buffer 20x, 94.5% H_2_O) (Invitrogen) at 150 V followed by Coomassie R-250 staining. We noted that no ABS3 dimerization was observed when performed using “dark blue” cathode buffer, whose components are proprietary.

## Notes

### Competing Interest Statement

The authors have declared no competing interest.

## Citations

Atherton, P. et al. (2015) Vinculin controls talin engagement with the actomyosin machinery, Nat. Commun., 6, pp. 1–12. doi: 10.1038/ncomms10038.

Azizi, L. et al. (2020) Cancer associated talin point mutations disorganise cell adhesion and migration, BioRxiv, pp. 1–28. doi: 10.1101/2020.03.25.008193.

Bate, N. et al. (2012) Talin contains a C-terminal calpain2 cleavage site important in focal adhesion dynamics, PLoS ONE, 7. pp. e34461. doi: 10.1371/journal.pone.0034461.

Buckley, C. D. et al. (2014) The minimal cadherin-catenin complex binds to actin filaments under force, Science, 346. 1254211. doi: 10.1126/science.1254211.

Cabantous, S., Terwilliger, T. C. and Waldo, G. S. (2005) Protein tagging and detection with engineered self-assembling fragments of green fluorescent protein, Nat. Biotech., 23, pp. 102–107. doi: 10.1038/nbt1044.

Franco-Cea, A. et al. (2010) Distinct developmental roles for direct and indirect talin-mediated linkage to actin, Dev. Biol., 345, pp. 64–77. doi: 10.1016/j.ydbio.2010.06.027.

Giannone, G. et al. (2003) Talin1 is critical for force-dependent reinforcement of initial integrin – cytoskeleton bonds but not tyrosine kinase activation, J. Cell. Biol., 163, pp. 409–419. doi: 10.1083/jcb.200302001.

Gingras, A. R. et al. (2005) Mapping and Consensus Sequence Identification for Multiple Vinculin Binding Sites within the Talin Rod, J. Biol. Chem., 280, pp. 37217–37224. doi: 10.1074/jbc.M508060200.

Gingras, A. R. et al. (2008) The structure of the C-terminal actin-binding domain of talin, EMBO J., 27, pp. 458–469. doi: 10.1007/s12104-007-9073-5.

Goult, B. T. et al. (2008) NMR assignment of the C-terminal actin-binding domain of talin, Biomol. NMR Assign., 2, pp. 17–19. doi: 10.1007/s12104-007-9073-5.

Goult, B. T., Yan, J. and Schwartz, M. A. (2018) Talin as a mechanosensitive signaling hub, J. Cell Biol., 217, pp. 3776–3784. doi: 10.1083/jcb.201808061.

Huang, D. L. et al. (2017) Vinculin forms a directionally asymmetric catch bond with F-actin, Science, 357, pp. 703–706. doi: 10.1126/science.aan2556.

Jiang, G. et al. (2003) Two-piconewton slip bond between fibronectin and the cytoskeleton depends on talin, Nature, 424, pp. 334–337. doi: 10.1038/nature01805.

Klapholz, B. et al. (2015) Alternative mechanisms for talin to mediate integrin function, Curr. Biol., 25, pp. 847–857. doi: 10.1016/j.cub.2015.01.043.

Klapholz, B. and Brown, N. H. (2017) Talin – the master of integrin adhesions, J. Cell Sci., 130, pp. 1–12. doi: 10.1242/jcs.190991.

Kopp, P. M. et al. (2010) Studies on the morphology and spreading of human endothelial cells define key inter- and intramolecular interactions for talin1, Eur. J. Cell. Biol., 89, pp. 661–673. doi: 10.1016/j.ejcb.2010.05.003.

Omabegho, T. et al. (2018) Controllable molecular motors engineered from myosin and RNA, Nat. Nanotech., 13, pp. 34–40. doi: 10.1038/s41565-017-0005-y.

Rahman, A. et al. (2016) Vinculin regulates directionality and cell polarity in two-And three-dimensional matrix and three-dimensional microtrack migration, Mol. Biol. Cell., 27, pp. 1431–1441. doi: 10.1091/mbc.E15-06-0432.

Ringer, P. et al. (2017) Multiplexing molecular tension sensors reveals piconewton force gradient across talin-1, Nat. Meth. 14, pp. 1090–1096. doi: 10.1038/nmeth.4431.

del Rio, A. et al. (2009) Stretching Single Talin Rod Molecules Activates Vinculin Binding, Science, 323, pp. 638–641. doi: 10.1126/science.1162912

Roca-Cusachs, P. et al. (2009) Clustering of alpha(5)beta(1) integrins determines adhesion strength whereas alpha(v)beta(3) and talin enable mechanotransduction, Proc. Nat. Acad. Sci. USA., 106, pp. 16245–50. doi: 10.1073/pnas.0902818106.

Schvartzman, M. et al. (2011) Nanolithographic control of the spatial organization of cellular adhesion receptors at the single-molecule level, Nano Lett., 11, pp. 1306–1312. doi: 10.1021/nl104378f.

Senetar, M. A., Foster, S. J. and McCann, R. O. (2004) Intrasteric inhibition mediates the interaction of the I/LWEQ module proteins Talin1, Talin2, Hip1, and Hip12 with actin, Biochemistry, 43, pp. 15418–15428. doi: 10.1021/bi0487239.

Smith, S. J. and McCann, R. O. (2007) A C-terminal dimerization motif is required for focal adhesion targeting of Talin1 and the interaction of the Talin1 I/LWEQ module with F-actin, Biochemistry, 46, pp. 10886–10898. doi: 10.1021/bi700637a.

Spudich, J. and Watt, S. (1971) The Regulation of Rabbit Skeletal Muscle The Regulation of Rabbit Skeletal Muscle Contraction, J. Biol. Chem., 246, pp. 4866–4871.

Srivastava, J. et al. (2008) Structural model and functional significance of pH-dependent talin-actin binding for focal adhesion remodeling, Proc. Nat. Acad. Sci. USA., 105, pp. 14436–14441. doi: 10.1073/pnas.0805163105.

Sung, J. et al. (2010) Single-Molecule Dual-Beam Optical Trap Analysis of Protein Structure and Function, Meth. Enzymol., 75, pp. 321–377.

Thievessen, I. et al. (2015) Vinculin is required for cell polarization, migration, and extracellular matrix remodeling in 3D collagen, FASEB J., 29, pp. 4555–4567. doi: 10.1096/fj.14-268235.

Yao, M. et al. (2014) Mechanical activation of vinculin binding to talin locks talin in an unfolded conformation, Sci. Rep., 4, p. 4610. doi: 10.1038/srep04610.

Yao, M. et al. (2016) The mechanical response of talin, Nat. Commun., 7, p. 11966. doi: 10.1038/ncomms11966.

Zhu, L., Yang, J., Bromberger, T. et al. (2017) Structure of Rap1b bound to talin reveals a pathway for triggering integrin activation. Nat. Commun., 8, 1744. doi: 10.1038/s41467-017-01822-8

